# Population genomics on the origin of lactase persistence in Europe and South Asia

**DOI:** 10.1101/2020.06.30.179432

**Authors:** Yoko Satta, Naoyuki Takahata

## Abstract

The C to T mutation at rs4988235 located upstream of the lactase (*LCT*) gene is the primary determinant for lactase persistence (LP) that is prevalent among Europeans and South Asians. Here, we review evolutionary studies of this mutation based on ancient and present-day human genomes with the following concluding remarks: the mutation arose in the Pontic Steppe somewhere between 23,000 and 5960 years ago, emigrated into Europe and South Asia in the Bronze Age via the expansion of the Steppe ancestry, and experienced local hard sweeps with their delayed onsets occurring between 5000 and 3280 years ago. We also argue that the G to A mutation at rs182549 arose earlier than 23,000 years ago, the intermediate CA haplotype ancestral to the LP-related TA haplotype is still represented by samples from Tuscans, admixed Americans and South Asians, and the great majority of G to A mutated descendants have hitchhiked since the C to T mutation was favored by local selection.

## Introduction

Most mammals lose the ability to digest lactose after weaning, but some present-day humans continue to express the key digestion enzyme, lactase (LCT), in the small intestine throughout adult life (Enattah et al., 2002; Ségurel and Bon, 2017). This physiological change is referred to as lactase persistence (LP), and has attracted much attention particularly from the viewpoint of gene-culture coevolution, human self-domestication or niche construction (e.g., Aoki, 1986; Itan et al., 2009, 2010; Gerbault et al., 2011, 2013; O’Brien and Laland, 2012). Apart from LP in Africa (Ingram et al., 2007; Tishkoff et al., 2007), LP in Eurasia is tightly associated with derived alleles T at rs4988235 and A at rs182549 of single nucleotide polymorphisms (SNPs) that are located -13,910 bp and -22,018 bp, respectively, upstream of the *LCT* gene. The association of the C/T polymorphism at rs4988235 with non-LP/LP is perfect in Finnish and non-Finnish (Korean, Italian and German) samples, and that of the G/A polymorphism at rs182549 is nearly perfect in case-control study samples (Enattah et al., 2002). Both the T and A alleles enhance the *LCT* promoter activity, although the A allele results in minimal enhancement compared with the T allele (Olds and Sibley, 2003).

Aside from the adaptive significance, the dating of LP or T allele origin in Europe has been of particular interest in light of ancient genomes. Using early Holocene human remains, Burger et al. (2007) showed that most Meso- and Neolithic Europeans lacked the T and A alleles, modestly concluding that LP arose in the last 20,000 years – hereafter denoted as *t*_*LCT*_ < 20,000 for convenience (Bersaglieri et al., 2004; Coelho et al., 2005; Leonardi et al., 2012). Lack of the T allele in European Neolithic farmers was directly demonstrated by the Chalcolithic Tyrolean Iceman, a 5300-year-old natural mummy discovered in the Ötztal Alps (Keller et al., 2012) as well as a 7400-year-old Cardial individual from Cova Bonica in Barcelona (Olalde et al., 2015). Obviously, however, more individuals were needed to confirm this absence. A recent study of 400 Neolithic, Chalcolithic and Bronze Age Europeans (Olalde et al., 2018) observed that the T allele remained at a very low frequency across the transition from the Neolithic period to the Bell Beaker and Bronze Age periods, both in Britain and continental Europe, with a major increase in its frequency occurring only within the last 3500 years (Cassidy et al., 2016; Brace et al., 2018). Table 1 summarizes some such results of Allentoft et al. (2015), Mathieson et al. (2015), and Olalde et al. (2018) based on the Central European chronology in Haak et al. (2015); readers may further refer to Witas et al. (2015), Liebert et al. (2017) and Ségurel and Bon (2017) together with Ségurel et al. (2020) and Jeong et al. (2020) both of which address an enigmatic 5000-year history of lactose adaptation in Central and Eastern Asian herders.

**Table 1.** Frequencies of the T allele at rs4988235 in ancient and present-day genomes in Central Europe.

| Central Europe | Cultures in Europe and Steppe | a | b | c |
| --- | --- | --- | --- | --- |
| Paleolithic<br>43,000-10,000 BC | Pleistocene hunter-gatherer | 0 | - | 0 |
| Mesolithic<br>9700-6200 BC | Mesolithic hunter-gatherer<br>8.2 event <sup>d</sup> | 0.17 <sup>e</sup> | 0 | 0 |
| Early Neolithic<br>6000-4000 BC | Holocene hunter-gatherer<br>LBK (5500 -4800BC) | 0 | 0 | 0 |
| Middle Neolithic<br>4000-3000 BC | Late Copper Age<br>Yamnaya (3000-2400 BC) | 0 | 0 | 0 |
| Late Neolithic<br>2600-2200 BC | CWC (2800-2300 BC)<br>BBC (2600-2000 BC) | ~0.20 | ~0.05 | ~0.05 <sup>f</sup> |
| Bronze Age<br>2200-1000 BC | Sintashta (2100-1800 BC)<br>Andronovo (1700-1500 BC) | 0.05 | ~0.05 | - |
| Iron Age 900 BC |  | - | - | ~0.05 <sup>f</sup> |
| Modern Europeans <sup>g</sup> |  | 0.67 | 0.74 | 0.46 |

The geographic location of the origin of European LP has been vague and contentious as well (Enattah et al., 2002, 2007 for the Pontic Steppe or Caucasian origin; Itan et al., 2009 and Gerbault et al., 2011 for theoretical modeling of LP/dairying coevolution) partly because the current geographic center of the LP distribution does not necessarily coincide with its location of origin when a population migrates and/or expands (Edmonds et al., 2004). Gerbault et al. (2011) provided an interpolated map of the distribution of the T allele in the Old World and summarized simulation results for the spread of European LP in time and space. Indeed, the T allele frequency varies considerably even within Europe (the 1000 Genomes Project Consortium, 2015; subsequently, we abbreviate the sequence dataset as 1000GD): 74% in Northern Europeans from Utah (CEU), 59% in Finnish from Finland (FIN), 72% in British from England and Scotland (GBR), 46% in Iberians from Spain (IBS), and 8.9% in Tuscans from Italy (TSI), with the overall frequency being 511/1006 = 51% in the European meta-population (EUR).

To gain some additional insights into these problems regarding LP or the T allele at rs4988235 and the A allele at rs182549, we applied our inference method, briefly described below, to an *LCT* region of 100 kb for the five individual European populations and one South Asian population (PJL: Punjabi from Lahore) in the 1000GD, separately (Smith et al., 2018). We then summarize and discuss results of the application together with the phylogenetic analysis of *LCT* haplotypes in light of recent ancient genomic studies. In addition, we demonstrate that one variable used in the method can pinpoint rs4988235 as the causative mutation for LP.

## Inference of a selective sweep under nonequilibrium demography

To detect, classify and date an incomplete selective sweep for *LCT* in each population, we used the method of the two-dimensional site frequency spectrum {*φ*_*i,j*_}, 2D SFS, by Satta et al. (2019). This method first seeks for a region under strong linkage disequilibrium (LD with *r*^2^ > 3/4) surrounding a target site and divides a sample of size *n* into the derived and ancestral allele groups defined by the site. Each element of the {*φ*_*i,j*_} matrix then counts the number of segregating sites or SNPs within the region at each of which we find exactly *i* and *j* derived alleles in the derived and ancestral allele groups, respectively. It is to be noted that *φ*_*i,j*_ > 0 for possible positive *i* and *j* stems most likely from recombination between the two allele groups, although it is expected to be rare in a tightly linked region.

We may compare the performance of our inference method with that of recent studies. Such studies include Itan et al. (2009) who estimated 6256 < *t*_*LCT*_ < 8683 using an approximate Bayesian computation (ABC) approach, Field et al. (2016) who developed the singleton density score method and estimated *t*_*LCT*_ ≥ 2000 from the UK10K project, Smith et al. (2018) who developed a hidden Markov model and obtained *t*_*LCT*_ = 9341 (95% CI: 8688-19,989) for IBS, 6869 (5143-8809) for BEB (Bengali in Bangladesh) and 9514 (8596-10,383) for PJL, Harris et al. (2018) who based their method on haplotype structure, Akbari et al. (2018) who claimed the high power of their iSAFE method to pinpoint the causative mutation of a single sweep, Nakagome et al. (2019) who developed an ABC method conditioned on the allele frequency in the past and applied it to four pigmentation SNPs, Satta et al. (2019) who used the 2D SFS method for EUR and estimated *t*_*LCT*_ = 3280 ± 480, and Tournebize et al. (2019) who obtained *t*_*LCT*_ = 4250 (95% CI: 3700-17,680) for CEU. Readers may also refer to Smith et al. (2018) and Tournebize et al. (2019) for various estimates of *t*_*LCT*_ either as the time to the most recent common ancestor (TMRCA) or as the age of the allele (Tishkoff et al., 2007; Nakagome et al., 2016).

Here, we focus on the TMRCA within the derived allele group because our interest is in the time (*t*_*SEL*_) of onset of positive selection rather than the origination time (*t*_*AGE*_) of a mutation that produced the allele. For convenience, we use the symbol *t*_*LCT*_ to denote the TMRCA of the derived allele group. Obviously, *t*_*AGE*_ is no shorter than *t*_*LCT*_, and *t*_*SEL*_ is no greater than *t*_*AGE*_, although none of these times is directly observable. For instance, to infer *t*_*SEL*_, Enattah et al. (2007) had to assume the initial frequency of the T allele and the selection intensity in a simple equation of describing an allele frequency change. Nevertheless, the distinction between *t*_*LCT*_ and *t*_*SEL*_ is important with respect to the mode of selective sweep. A necessary condition for *soft* sweeps (Hermisson and Pennings, 2005, 2017) is *t*_*LCT*_ > *t*_*SEL*_. This is equivalent, though not exactly, to saying that multiple distinct ancestral lineages carrying a common mutation simultaneously undergo positive selection. An obvious caution is that positive selection acting on a standing variation does not necessarily classify the sweep as soft. To the contrary, a sufficient condition for *hard* sweeps is *t*_*LCT*_ ≤ *t*_*SEL*_. There are thus two distinguishable cases: hard (*t*_*LCT*_ ≤ *t*_*SEL*_ ≤ *t*_*AGE*_) and *conditional* soft sweeps (*t*_*SEL*_ < *t*_*LCT*_ ≤ *t*_*AGE*_), the latter being related to the hardening of soft sweeps (Wilson et al., 2014). It seems that either origin of a target mutation or population structure is irrelevant or even confusing to this distinction.

To evaluate statistical significances of our estimates, we assumed a demographic model and neutrality of genome evolution (Kimura, 1968). We used a very simplified bottleneck and expansion model that incorporates some aspects of human nonequilibrium demographic history inferred thus far by SFS (Schaffner et al., 2005), PSMC (Li and Durbin, 2011), MSMC (Schiffels and Durbin, 2014), Stairway Plot (Liu and Fu, 2015), and SMC++ (Terhorst et al., 2017) among others. In our simplified model, a single bottleneck is shared by all non-African populations from the exodus out of Africa (ca. 58,000 years ago) to the initial Eurasia-wide dispersal (ca. 45,000 years ago), and a population had since grown exponentially up to the population size 500 years ago (Figure 1, Table 2a). We assumed the size of the bottlenecked population to be 2000 and ignored the effect of population explosion during the last 500 years. The robustness of the method to nonequilibrium demography was discussed in Satta et al. (2019). For genome evolution, we assumed the infinite-sites model (Kimura, 1969), but with linkage.

**Table 2a.** IAV variables in the 100 kb LD region surrounding rs4988235 in five European populations in the 1000GD (sample size n and derived allele frequency f_r_ = m/n).

| Population | CEU | FIN | GBR | IBS | TSI |
| --- | --- | --- | --- | --- | --- |
| $\alpha$ | $1.3 \times 10^{-4}$ | $3.6 \times 10^{-5}$ | $5.1 \times 10^{-5}$ | $6.0 \times 10^{-5}$ | $5.1 \times 10^{-5}$ |
| $m/n$ | 146/198 | 117/198 | 131/182 | 98/214 | 19/214 |
| $S_\xi$ | 213 | 218 | 204 | 287 | 351 |
| $F_c$ | 0.0141 | 0.0076 | 0.0135 | 0.0017 | 0.0059 |
| ($P$) | (< $10^{-4}$) | (< $10^{-4}$) | (< $10^{-4}$) | (< $10^{-4}$) | (0.015) |
| $L_{c0}$ | 0.0026 | 0.0021 | 0.0026 | 0.0003 | 0.0002 |
| ($P$) | (< $10^{-4}$) | (< $10^{-4}$) | (< $10^{-4}$) | (< $10^{-4}$) | (0.013) |
| $G_{c0}$ | 1.09 | 1.25 | 1.16 | 1.50 | 1.00 |
| ($P$) | (< $10^{-4}$) | (< $10^{-4}$) | (< $10^{-4}$) | (< $10^{-4}$) | (0.036) |
| $C$ | 0.164 | 0.171 | 0.168 | 0.184 | 0.105 |
| (S. E.) | (0.036) | (0.047) | (0.042) | (0.066) | (0.074) |

The $P$ values in parentheses were computed from simulation results of $10^4$ replications of $ms$ (Hudson, 2002) under the bottleneck and exponential growth model
with the per-year rate $\alpha$ (Figure 1). The variables are defined as $F_c = \frac{\sum i\varphi_{i,j}}{\sum (i+j)\varphi_{i,j}}$ ,
$L_{c0} = L_{\zeta 0}/L_\xi$ , $G_{c0} = L_{\zeta 0}/S_{\zeta 0}$ and $C = L_{\zeta 0}/m$ where $\Sigma$ in $F_c$ indicates that the summation is taken over a certain range of $i$ and $j$ (see Fujito et al., 2018 and Satta et al., 2019 for details), $S_{\zeta 0} = \sum_{i=1}^{m-1} \varphi_{i,0}$ , $L_{\zeta 0} = \sum_{i=1}^{m-1} i\varphi_{i,0}$ , $S_\xi = \sum_{i=0}^m \sum_{j=0}^{n-m} \varphi_{i,j}$ , and $L_\xi = \sum_{i=0}^m \sum_{j=0}^{n-m} (i+j)\varphi_{i,j}$ .

**Figure 1.**
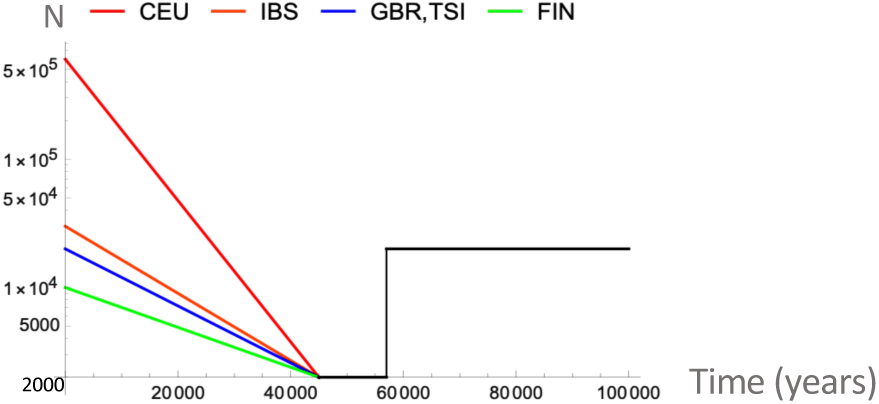
Demographic models for five European populations. The effective population size (Y axis in log scale) is depicted against the past time (X axis in years). Four colored lines show population-specific demographic models with different exponential growth rates (*α*) (Table 1). The models assume that the size (500 years ago) was 600,000 for CEU, 30,000 for IBS, 20,000 for GBR and TSI, and 10,000 for FIN. All populations began to expand 45,000 years ago after experiencing a bottleneck phase of constant size 2000 since this ancestral Eurasian population migrated out of Africa 58,000 years ago. The ancestral African population size was 20,000.

## Results

The hallmark of an incomplete selective sweep is reduced intra-allelic variability (IAV) within a derived allele group relative to the total variability in the entire sample (Fujito et al., 2018; Satta et al., 2019). IAV may be expressed by numerous variables that are useful for detecting and classifying a selective sweep as well as for dating the onset of positive selection. The significance test for individual variables *F*_*c*_, *L*_*c*0_ and *G*_*c*0_ used to measure the level and pattern of IAV shows that the observed values are as extreme as 10^−4^ in all but the TSI population (Table 2a). To further strengthen the test, we may combine these mutually correlated variables, but to avoid inflating false positives, we used the method by Brown (1975) that combines non-independent, one-sided tests of significance (Poole et al., 2016). The results of this test show that, in the European populations including TSI, the individual IAVs become highly incompatible with neutrality (Table 2b).

**Table 2b.**
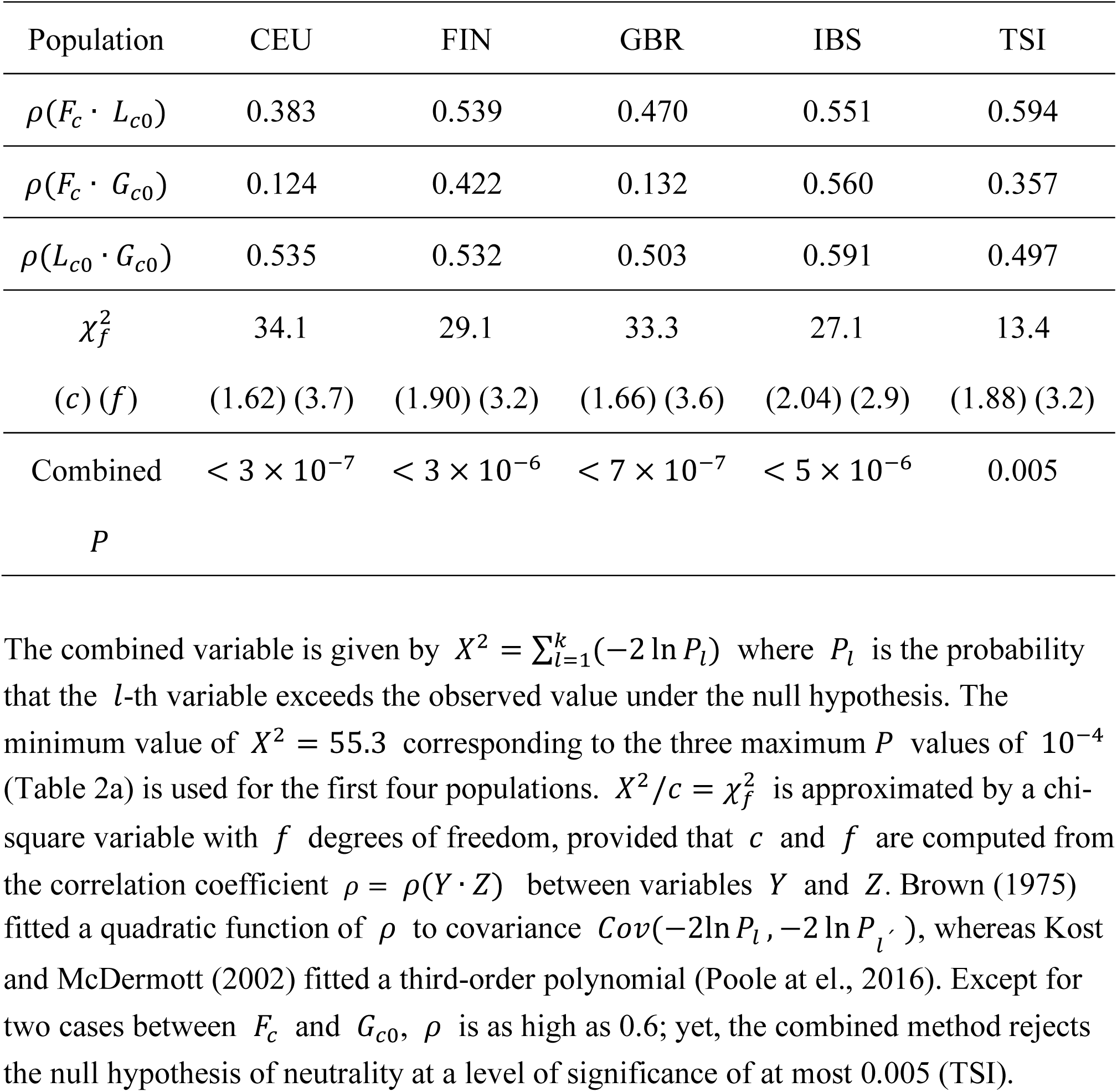
P values obtained by the method that combines non-independent, one-sided F_c_, L_c0_ and G_c0_ variables.

The 2D SFS method then infers that the selective sweep for the T allele is definitely hard. In addition to unusually reduced IAV levels measured by *F*_*c*_ and *L*_*c*0_, this hardness is captured as low levels of multiplicity (*G*_*c*0_) of derived alleles per SNP (Table 2a). Indeed, almost all alleles per SNP are singletons or doubletons, irrespective of populations, such that *G*_*c*0_ is close to 1 to 2. No other target sites that exhibit such low multiplicity in such a wide genomic region have yet to be revealed (Satta et al., 2019). The 2D SFS method also allows us to estimate the mutation-rate-scaled TMRCA (*ut*_*LCT*_) from the total number 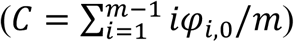 of derived alleles per sample within the T allele group of size *m* < *n*. It is to be noted that recombinant SNP sites are automatically excluded from *C*. Using *ut*_*LCT*_ = *C* or *t*_*LCT*_ = *C*/*u* (Table 2a), we have *t*_*LCT*_ = 3280 ± 720 in CEU of *m* = 146, 3420 ± 940 in FIN of *m* = 117, 3360 ± 840 in GBR of *m* = 131, 3380 ± 1320 in IBS of *m* = 98, and 2100 ± 1480 in TSI of *m* = 19, if the above mutation rate (*u*) is 5 × 10^−5^ per 100 kb per year (Scally and Durbin, 2012).

We also conducted phylogenetic analysis to show that almost all T haplotypes (haplotypes within the T allele group defined in the LD region), including those in TSI and PJL, are tightly clustered so as to form a single clade (Figure 2). Unexpectedly, there is one T haplotype (HG03653) in PJL that is clustered in the ancestral C allele group. The DNA sequence of HG03653 suggests three possibilities: a mistyping of C to T at rs4988235, a parallel mutation in a divergent haplotype background resulting in the TG haplotype, or a migrant of the TG haplotype from the Urals or the Caucasus as discussed below. In general and more importantly, the ancestral C haplotype closest to the T cluster differs only by a single C to T mutation. This confirms the previous conclusion that the T haplotypes were descended from an ancestral C haplotype in the recent past by a single mutation (Enattah et al., 2007).

**Figure 2.**
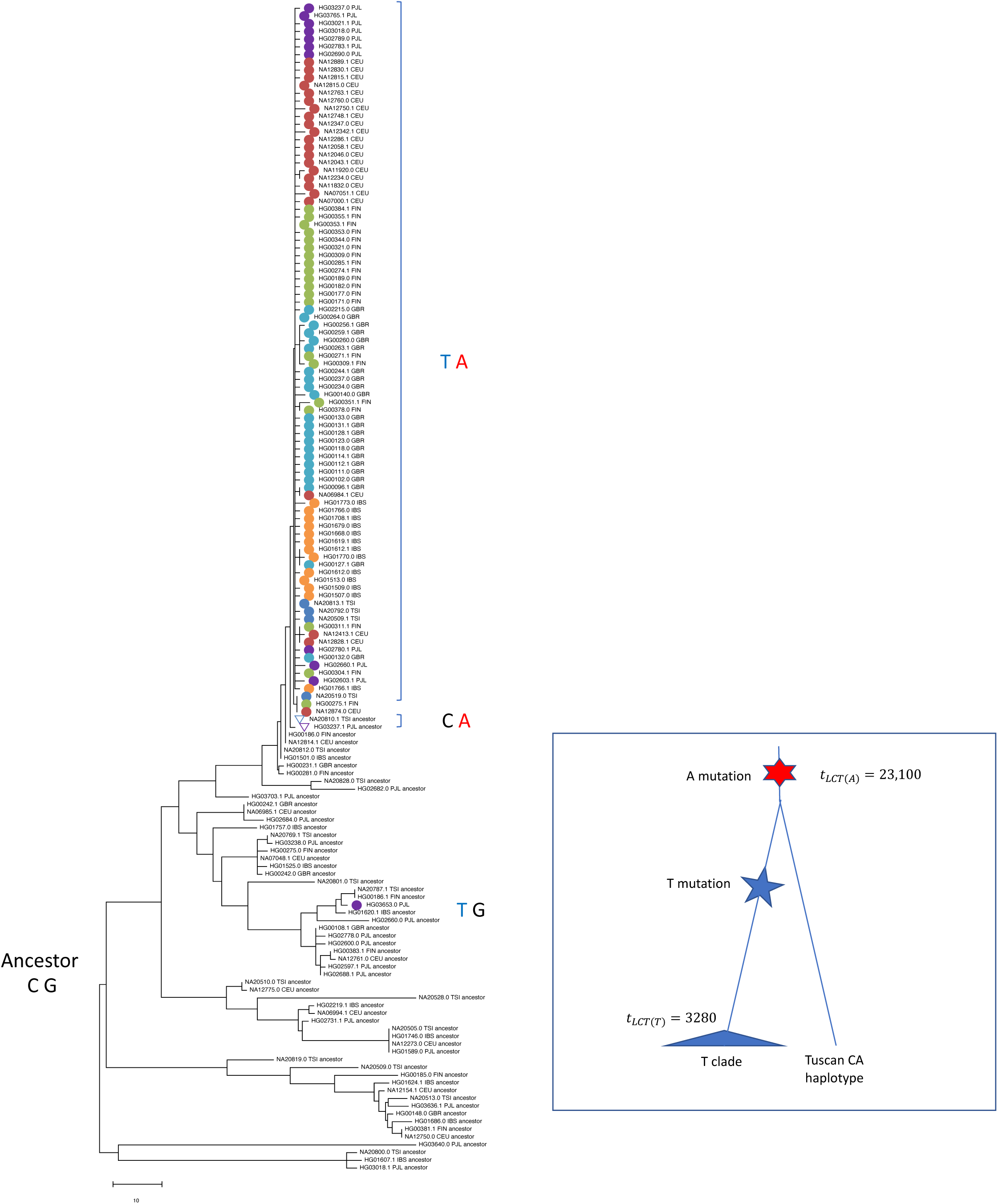
Unrooted NJ tree for a subsample of the T and C distinct haplotypes. The T/C sample size is 23/9 from CEU, 13/9 from IBS, 22/5 from GBR, 4/12 from TSI, 20/7 from FIN, and 11/15 from PJL. The LD region in each population is the same as that in the entire EUR sample (Satta et al., 2019). TA haplotypes at rs4988235 and rs182549 are marked by filled colored circles and CA haplotypes in TSI and PJL by two open inverse triangles. The tree was constructed from pairwise nucleotide differences without multiple-hit correction (Saitou and Nei, 1987). The inset is a cartoon of the haplotype evolution. The G to A mutation occurred >23,000 years ago, the C to T mutation emerged much later in an A lineage and swept through a population. The CA haplotype in TSI and PJL is a remnant of the A mutation that escaped from the selective sweep.

It is also interesting to examine how accurately some of our variables can pinpoint a true target site of positive selection (Figure 3). For this purpose, we selected *F*_*c*_ based on the fact that recombination plays essential roles in pinpointing a target site and that *F*_*c*_ includes recombinant SNPs, whereas both *L*_*c*0_ and *G*_*c*0_ do not. Nominating all candidate SNPs in a 2.5 Mb genomic region surrounding *LCT*, we measured *F*_*c*_ for each candidate within a window of 300 SNPs and then ranked all of these *F*_*c*_ values in the whole 2.5 Mb region. We also compared the result with that by the iSAFE with an “inserted” procedure (Akbari et al., 2018), and found that this procedure does not improve the power of *F*_*c*_. Most importantly, like iSAFE, *F*_*c*_ ranked the SNP at rs4988235 as 1.

**Figure 3.**
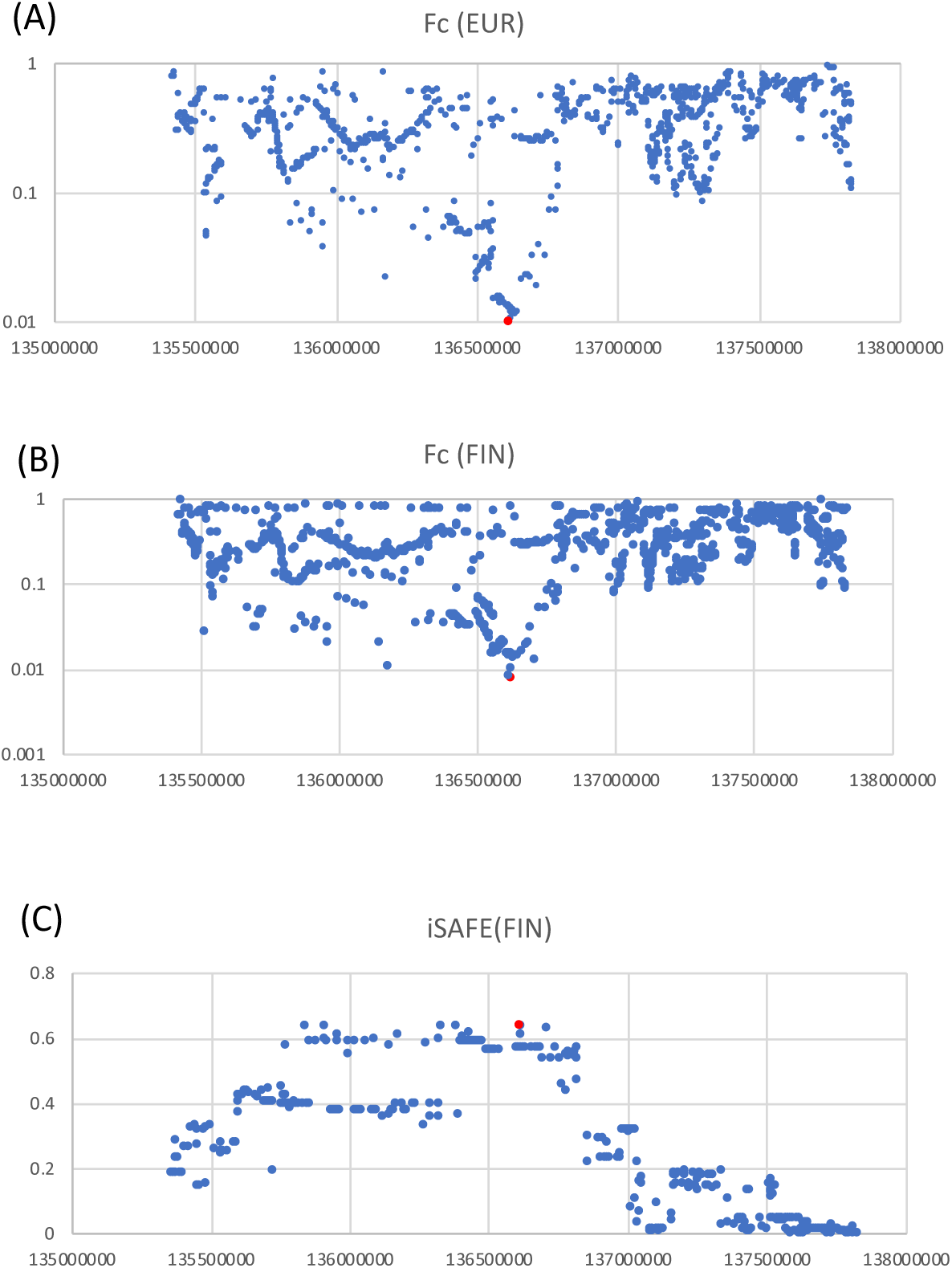
Identification of a target site of selection. We searched for a target site from candidate SNPs surrounding *LCT* (X axis for genomic positions) by ranking their *F*_*c*_ and iSAFE values (Y axis). In *F*_*c*_, candidate SNPs were all derived alleles whose frequencies are >45% in EUR and >54% in FIN. Note that, although derived alleles at low frequencies were excluded from candidates, they were used to compute *F*_*c*_. The red points indicate the *F*_*c*_ or iSAFE value for the T allele at rs4988235. Needless to say, the ranking by *F*_*c*_ is in ascending order (Fujito et al., 2018), whereas that of iSAFE is in descending order (Akbari et al., 2018). (A), (B) *F*_*c*_ scanning over the 2.5 MB genomic region in the EUR sample of size *n* = 1006 and the FIN sample of size *n* = 189; (C) iSAFE scanning over the 2.5 MB genomic region in the FIN sample.

## Discussion

Farming was introduced to most of Europe from Anatolia/Levant by 6000 to 5000 BC via two main routes: the Danube river line and the northern Mediterranean coast line. Farmers in Central Europe and Iberia were respectively associated with the LBK (Linearbandkeramik) tradition along the river line and the *Impresso*-Cardium culture along the coast line (Olalde et al., 2015; Hofmanová et al., 2016; Broushaki et al., 2016; Brace et al., 2018; Mathieson et al., 2018; Souilmi et al., 2020). In each route, farming was brought upon by demic diffusion with limited admixture from local hunter-gatherers and accordingly an apparent decrease in the Neolithic contribution with geographic distance from the Near East (Bramanti et al., 2009; Skoglund et al., 2014; Günther et al., 2015). Indeed, while Scandinavian farmers were intimately related to farmers in Southern Europe, such as the Tyrolean Iceman as well as Sardinians and many other present-day groups, they exhibited high levels of admixture from local hunter-gatherers (Skoglund et al., 2012, 2014). Rivollat et al. (2020) also revealed a higher proportion of western hunter-gatherer ancestry in Western European farmers than in Central European farmers, pointing to the complexity of their interactions in France where both routes converged.

Olalde et al. (2018) showed that Neolithic Anatolian and Aegean farmers were indeed non-LP despite archaeological and genetic evidence for their taurine cattle domestication and milk use around 6500 BC (Evershed et al., 2008; Curry, 2013). Allentoft et al. (2015) found the T allele at low frequency in the Bronze Age of Eurasia (ca. 3000-1000 BC) and showed a possible Steppe origin of LP from the highest occurrence of the T allele in the Yamnaya and Corded Ware cultures (CWCs) between 3000/2800 and 2300/2000 BC. Haak et al. (2015) also observed that the T allele increased in frequency only after the Yamnaya Steppe ancestry became ubiquitous in Central and Western Europe. Consistent with these findings, it was recently identified that the T allele is possessed by only one CWC individual from Sweden (Malmström et al., 2019), only one Final Neolithic individual from Switzerland (Furtwängler et al., 2020), and none of 16 CWC individuals from Poland (Witas et al., 2012; Linderholm et al., 2020), to mention a few. Using 230 ancient Eurasians, Mathieson et al. (2015) found that the earliest appearance of the T allele is in a Central European Bell Beaker sample dated to between 2450 and 2140 BC, and the selective sweep dated to the last 4000 years (Burger et al., 2007; Gamba et al., 2014; Haber et al., 2016 for a review). Bell Beaker culture (BBC) was widespread in Western and Central Europe from 2750/2500 to 2200/1800 BC, and associated with Steppe-related ancestry with the replacement in gene pool being most pronounced in Britain (Olalde et al., 2018). Interestingly, however, Northern Italy and Sicily Bell Beakers had no sign of LP. Raveane et al. (2019) and Fernandes et al. (2020) pointed out that they can be modeled certainly as Anatolian-farmer-related, though with different affinities to Steppe Bronze Age. Olalde et al. (2018) also argued that there is limited genetic affinity between Beaker-complex-associated individuals from Iberia and Central Europe. Furthermore, focusing on the 8000-year history of Iberia, Olalde et al. (2019) concluded that, in Iberia, the T allele continued to occur at low frequency in the Iron Age, pointing to recent strong positive selection; however, they also showed that present-day non-Indo-European Basques inherited substantial levels of Steppe ancestry, consistent with Late Neolithic Basques with 27% LP (Plantinga et al., 2012). In short, although estimates of the ancient T allele frequency still vary, neither hunter-gatherers nor early farmers in Anatolia and Europe possessed the T allele (Table 1) and the earliest carriers came from CWC and BBC individuals in Central Europe, Iberia and Scandinavia after ca. 2500 BC.

It is therefore reasonable to assume that the T allele arose somewhere in the Pontic Steppe and emigrated into Central Europe via the Yamnaya expansion in ca. 3000 BC. Yet, positive selection began to operate for the immigrated T allele somewhat later. Literally speaking, as positive selection acted on such a standing variation (*t*_*SEL*_ < *t*_*AGE*_), the selective sweep might be defined as soft. Nonetheless, the level and pattern of IAV in the T allele group are best described as hard. In this context, it is not really important to distinguish between hard and soft sweeps based on the origin of a selected mutation. What can be done is to practically classify the mode based on IAV in a linked genomic region. The delayed onset of positive selection for European LP suggests the initial operation on genetically indistinguishable T haplotypes (*t*_*LCT*_ ≤ *t*_*SEL*_). This may result from either where selection operated on a single ancestral lineage or where multiple descendants, if involved, did not have enough time to differentiate from each other by mutations. In any event, both cases lead to *t*_*LCT*_ ≤ *t*_*SEL*_ < *t*_*AGE*_ and look like hard sweeps. Evaluation of *t*_*AGE*_ may be directly made from the age estimate of the T allele, but here we used the TMRCA (*t*_*LCT*(*A*)_) of the derived A allele group defined at rs182549 as a surrogate of the age of the T allele (*t*_*AGE*(*T*)_). The widespread T allele occurred in the background of the A allele and *t*_*AGE*(*T*)_ is bounded by *t*_*LCT*(*A*)_ = 23,100 ± 20,000 in EUR. Thus, *t*_*AGE*(*T*)_ must be somewhere between *t*_*LCT*(*T*)_ = 3280 and *t*_*LCT*(*A*)_ = 23,100: Ségurel et al. (2020) discovered that, in the currently available ancient genomes, the earliest appearance was in Ukraine 5960 years ago (Mathieson et al., 2018). As *t*_*LCT*(*A*)_ is well after non-African populations differentiated from each other, it is likely that both the T and A alleles originated in a single locally differentiated Eurasian population. However, there is an exception to this conclusion. From genotyping a ∼30 kb *LCT* region in >1600 samples from 37 global populations, Enattah et al. (2007) found that the T mutation occurred twice independently on highly divergent haplotype backgrounds and that one mutation is geographically restricted to populations living in west of the Urals and north of the Caucasus.

An interesting caveat is related to Tuscans. Clearly, the present-day TSI is an outlier in terms of European LP, exhibiting an exceptionally low T allele frequency in a large sample. By comparing TSI genomes with other Europeans and Middle Easterners, Pardo-Seco et al. (2014) revealed that admixture took place between 1100 and 600 BC. Although this admixture event may have been much older and even recurrent, it implies the partial eastern origin of the non-Indo-European Etruscans, ancestral to Tuscans in the Bronze/Iron Age. Intriguingly, some cattle breeds (*Bos taurus*) were also simultaneously imported from the east Mediterranean area (Pellecchia et al., 2007). However, the T allele in TSI suggests that this cattle import did not act as a selective agency for LP due presumably to lack of relevant culture or absence of the T allele *per se* at that time. We emphasize that, although evidence for positive selection is relatively weak for TSI, the Brown’s combining method could detect a hard sweep for the T allele group (Table 2b). One possibility is that the T allele in TSI resulted from recent gene flow from neighboring LP areas (Fiorito et al., 2015 for gene flow within Italy). Coelho et al. (2005) estimated the LP frequency to be 24% and the T allele frequency to be 13% in Central Italy (37 individuals from Tocco da Causaria and 30 from Rome). While these frequencies in Central Italy are much lower than those in Portugal, they are much higher than those in Tuscany; if statistically tested, Central Italian genomes would undoubtedly show a hard sweep.

TSI is exceptional also in terms of pairwise LD between C/T at rs4988235 and G/A at rs182549. While the TA haplotype at these SNPs is in complete LD (*r*^2^ = 1) in CEU, FIN, GBR, and IBS, it is not (*r*^2^ = 0.95) in TSI, owing to the presence of a CA haplotype in addition to derived TA and ancestral CG haplotypes in the 1000GD (Figure 2). As a number of CA haplotypes can be also found in MXL (Mexican Ancestry from Los Angeles) and PEL (Peruvian in Lima) from admixed Americans (AMR), as well as PJL, BEB, ITU (Indian Telugu in the UK), and STU (Sri Lankan Tamil from the UK) from South Asian Ancestry (SAS), the age of the A allele must be older than that of the T allele as aforementioned. Interestingly, the T allele frequency is quite high in some of these populations: 24% in MXL, 11% in PEL, and 26% in PJL. Yet, IAV is substantially reduced in MXL or even non-existent in PEL (i.e., *t*_*LCT*_ = 0). These features are compatible with very recent ongoing selective sweeps, but a more likely cause would be admixture, founder effects, or a combination of these factors after Columbus. Similarly, IAV in PJL is significantly reduced (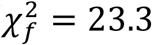 with *c* = 2.01 and *f* = 2.98; *P* = 4 × 10^LW^) with *t*_*LCT*_ = 3600 ± 1320 (Figure 2). Although admixture also occurred in PJL, we consider a prehistoric event as a more likely cause: if the T allele arose in the Pontic Steppe, it must have spread not only into Europe from 3000 BC, but also into Turan by 2100 BC and further into South Asia after the decline of the Indus Valley Civilization in ca. 1800 BC (Damgaard et al., 2018; Narasimhan et al., 2019; Shinde et al., 2019). As the time elapsed in Punjab was as long as 3800 years, it is tempting to speculate the presence of an independent selective sweep by the T allele in this locality too. Thus, it appears that LP in Europe and South Asia shares the early history of the expanding Steppe ancestry in the Bronze Age, and offers strong selective advantages to its local bearers in lactose-relevant niche construction.

## Acknowledgements

We thank Drs. Laure Ségurel, Seiji Kadowaki, Quintin Lau and Kenichi Aoki for their interest and comments. This work was supported in part by the Scientific Research on Innovative Areas, a MEXT Grant-in-Aid Project FY2016-2020.

